# Synergistic role of *Alu* and core duplicon sequences in driving genomic instability at the disease-associated 16p12.3–p13.11 region

**DOI:** 10.1101/2025.07.09.663898

**Authors:** Stefania Fornezza, Giuliana Giannuzzi

## Abstract

Human chromosome 16p is a hotspot of interspersed segmental duplications (SDs), which have served as row material for gene innovation while also increasing susceptibility to recurrent pathogenic rearrangements. These SDs are organized around a high-copy, ancestral core duplicon known as LCR16a (Low Copy Repeat 16a), and are characterized by an enrichment of *Alu* elements at their boundaries. In this study, we investigated the molecular mechanisms underlying SD formation across four clusters within the 16p12.3–p13.11 region. Analysis of 94 genome assemblies showed 12 and 8 alternative structures in clusters 1-2 and 3-4, respectively. These include at least 10 duplication/deletion events and 5 inversions. Breakpoint mapping revealed that 13 out of 15 events arose via nonallelic homologous recombination (NAHR). In particular, 6 had breakpoints in duplicons associated with LCR16a, 2 occurred within the LCR16a sequence itself, and 5 were mediated by *Alu* elements—one of which located within LCR16a. The last mechanism contributes to multiallelic, tandem copy-number variation involving two distinct LCR16a-duplicon pairs. We identified an interspersed, inverted duplicative transposition, possibly driven by double template switches at homologous sites— specifically, *Alu* elements located within inverted copies of LCR16a and an associated duplicon. Comparative analyses using nonhuman primate T2T genome assemblies suggest that clusters 3 and 4 originated ∼6 million years ago in the human-chimpanzee common ancestor through double template switching involving *Alu* repeats. Clusters 1 and 2 primarily formed more recently through human-specific LCR16a-associated duplicative transpositions. These findings highlight a synergistic interplay between *Alu* elements and LCR16a in driving 16p12.3–p13.11 SD formation and genomic instability, through both recombination-and replication-based mechanisms.

## INTRODUCTION

The short arm of human chromosome 16 is a hotspot for diverse copy-number variations associated with neurodevelopmental disorders (1–13). These recurrent chromosomal rearrangements arise from nonallelic homologous recombination (NAHR) between interspersed, highly identical segmental duplications (SDs), also known as low-copy repeats (LCRs) (1, 14). While SDs with ≥90% sequence identity and ≥1 kb in length account for approximately 7% of the human genome (15), their proportion increases to 11% on chromosome 16 and reaches 23% on its short arm, based on the T2T-CHM13v2.0 human genome assembly.

Comparative analyses have revealed an expansion of SDs in the common ancestor of great apes, along with a higher number of interspersed SDs in African great apes (16). This expansion is believed to be driven by recombination-based mechanisms mediated by high-copy-number repeats, such as *Alu* elements in the human genome (17, 18) and SVA (SINE-R-VNTR-*Alu*) elements in the orangutan genome (19). *Alu* elements are primate-specific 300 bp retrotransposons that are categorized into three subfamilies, each active at different times during primate evolution: *AluJ* (65–40 Mya, million years ago), *AluS* (45–25 Mya), and *AluY* (from 30 Mya to present) (20).

Human SD clusters are characterized by the presence of a “core duplicon”—an ancestral, high-copy segment shared by the majority of a group of intrachromosomal SDs—which appears to have driven the formation of interspersed duplications (21–23). For instance, the duplication architecture of chromosome 16p is centred around a 20 kbp segment known as LCR16a (Low Copy Repeat 16a), which embeds the *NPIP* (nuclear pore interacting protein) gene family (21, 24–26). LCR16a exists as a single copy in Old World monkeys (e.g., macaque and baboon genomes) but has undergone independent and recurrent expansions via duplicative transposition in various primate lineages, including humans, chimpanzees, gorillas, orangutans, and marmosets (25, 26).

NAHR and unequal crossing over between pairs of common or low-copy repeats account for the formation of SDs in a tandem configuration and are the primary mutational mechanisms responsible for recurrent copy-number variation (CNV) in humans (27). In contrast, the mechanisms underlying the emergence of novel interspersed SDs remain less well understood. A study of ∼1000 large structural variants in humans showed that the majority arose via microhomology-mediated processes or NAHR (28). More recently, some studies have suggested that replicative circular DNA intermediates may contribute to the generation of certain large SDs in the human genome (29, 30). Moreover, while the enrichment of *Alu* elements at SD edges and the presence of core duplicons are well established, their precise roles in SD formation, as well as the possible interplay between *Alu* sequences and core duplicons, remain unresolved.

In this study, we aimed to shed light on the mechanisms of SD formation and the contribution of *Alu* elements and core duplicons to these processes by analysing ongoing and evolutionary structural variation at the highly mutagenic and SD-rich 16p12.3–p13.11 locus. This region contains the ancestral copy of LCR16a (26) and is a hotspot for recurrent pathogenic copy-number variations, with breakpoints occurring within SD clusters (4, 5, 10, 31) (**Figure 1**).

**Figure 1.**
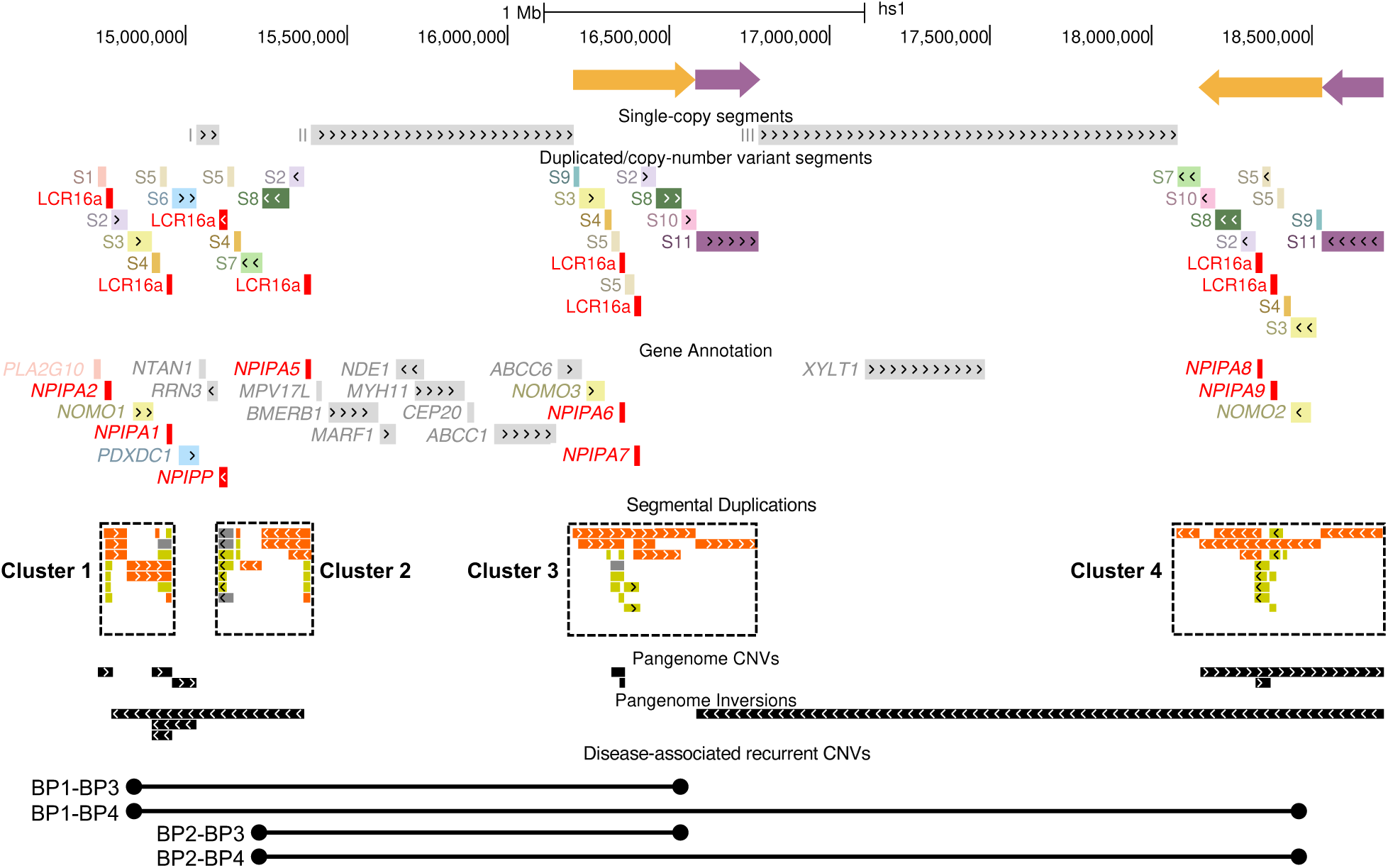
Genomic architecture of the human 16p12.3–p13.11 chromosomal region. View of the region Chr16:14,000,000-19,000,000 on human T2T-CHM13 v2.0 genome in the UCSC Genome Browser. Single-copy regions are illustrated as I, II, and III gray segments. We operationally divided segmental duplications (SDs) and copy-number variant intervals in 12 segments (S1 to S11, and LCR16a). Segments in clusters 3 and 4 correspond to two large, inverted SDs (yellow and purple arrows at the top). LCR16a core duplicons are shown in red. Protein-coding genes are color-coded according to the segment where they map. SDs are organized into four clusters (dashed boxes), with colors reflecting the similarity levels: 90–98% in light and dark gray; 98–99% in yellow; >99% in orange (52, 57). Copy-number variations (CNVs) and inversion polymorphisms identified in the pangenome dataset are shown with black segments. At the bottom, the intervals of pathogenic recurrent CNVs with breakpoints (BP) within SD clusters are shown.

## RESULTS

### 16p12.3–p13.11 structural diversity

SDs make up 36% of the chr16:14,000,000–19,000,000 interval (data according to T2T-CHM13 v2.0 reference), where they are organized into four clusters arranged around the LCR16a core duplicon (**Figure 1**). In the T2T-CHM13 genome, there are eight LCR16a copies at 16p12.3–p13.11, including seven *NPIPA* protein-coding genes and the noncoding *NPIPP* (**Figure 1**).

We operationally divided the region into three single-copy intervals of 70 kbp, 800 kbp, and 1.3 Mbp (indicated as I, II, and III in **Figure 1)** and 12 segments (S1 to S11 and LCR16a). All segments except S1 and S6 map within annotated SDs (**Figure 1**). The majority of high-identity SDs have paralogous copies in the four clusters, suggestive of recent duplications within the region. With the exception of segment S7 that corresponds to an SD pair between clusters 1 and 4, clusters 3 and 4 contain the same segments but in different configurations: the interval spanning segments S9 to S10 (yellow arrow in **Figure 1**) and segment S11 (purple arrow) are switched in position and inverted between clusters 3 and 4.

We analyzed this locus in 47 diploid, phased assemblies (94 genomes) from the Human Pangenome Project (32), representing an unselected sample to measure the extent of structural variation at this region. The entire interval was assembled in a single contig in 20 genomes (21%), highlighting challenges in its complete reconstruction. To improve sequence recovery, we divided the locus into two subintervals: the first one including clusters 1 and 2 (chr16:14,000,000-16,000,000) and the second one including clusters 3 and 4 (chr16:16,000,000-19,000,000). This approach yielded 38 contiguous sequences for clusters 1 and 2 (40%) and 34 for clusters 3 and 4 (37%), with a total of 39 and 35 sequences including those from the T2T-CHM13 genome (**Supplemental Table 1**).

We identified 12 alternative structures (H1–H12) within the clusters 1-2 subinterval. Structural variation consisted in three types of copy-number variations: i) tandem duplications of segments S1-LCR16a (tdup1) at the start of cluster 1 in seven haplotypes; ii) deletion of segments LCR16a-S5 (del1) in H3; and iii) deletion of LCR16a-S6-S4 (del2) in H8. We also identified three inversion polymorphisms (inv1, inv2, and inv3) (**Figures 1, 2, Table 1**). The direction of each rearrangement (e.g., deletion in H3 *versus* H2 instead of duplication in H2 *versus* H3) is based on our ordering of the 12 structures. We arranged them starting from the most frequent haplotype and linking subsequent structures along the most parsimonious trajectory (i.e., involving the fewest rearrangements), though this may not necessarily reflect the actual evolutionary order (**Figure 2**).

**Figure 2.**
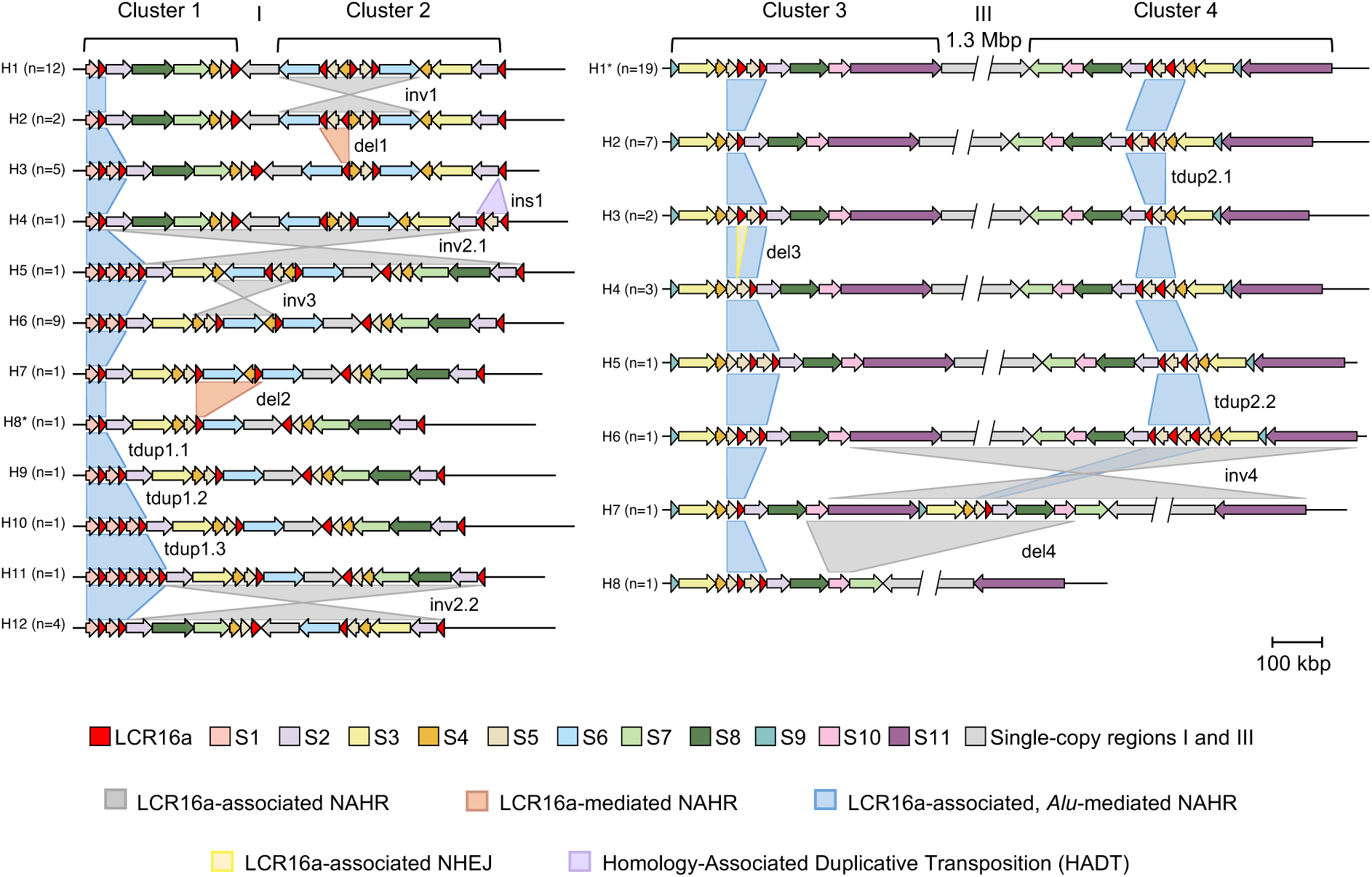
16p12.3–p13.11 structural diversity. Schematic representation of 12 and 8 different haplotypes of clusters 1-2 (*left*) and clusters 3-4 (*right*), identified in 53 haploid genomes. An asterisk designates the structures observed in the T2T-CHM13 reference genome (H8 for clusters 1-2; H1 for clusters 3-4). Colored links between haplotypes illustrate the structural variation and its mechanism of origin. The insertion “ins1” in cluster 2 is unique to H4; the deletion “del3” in cluster 3 is found in H4 and H5.

**Table 1.**
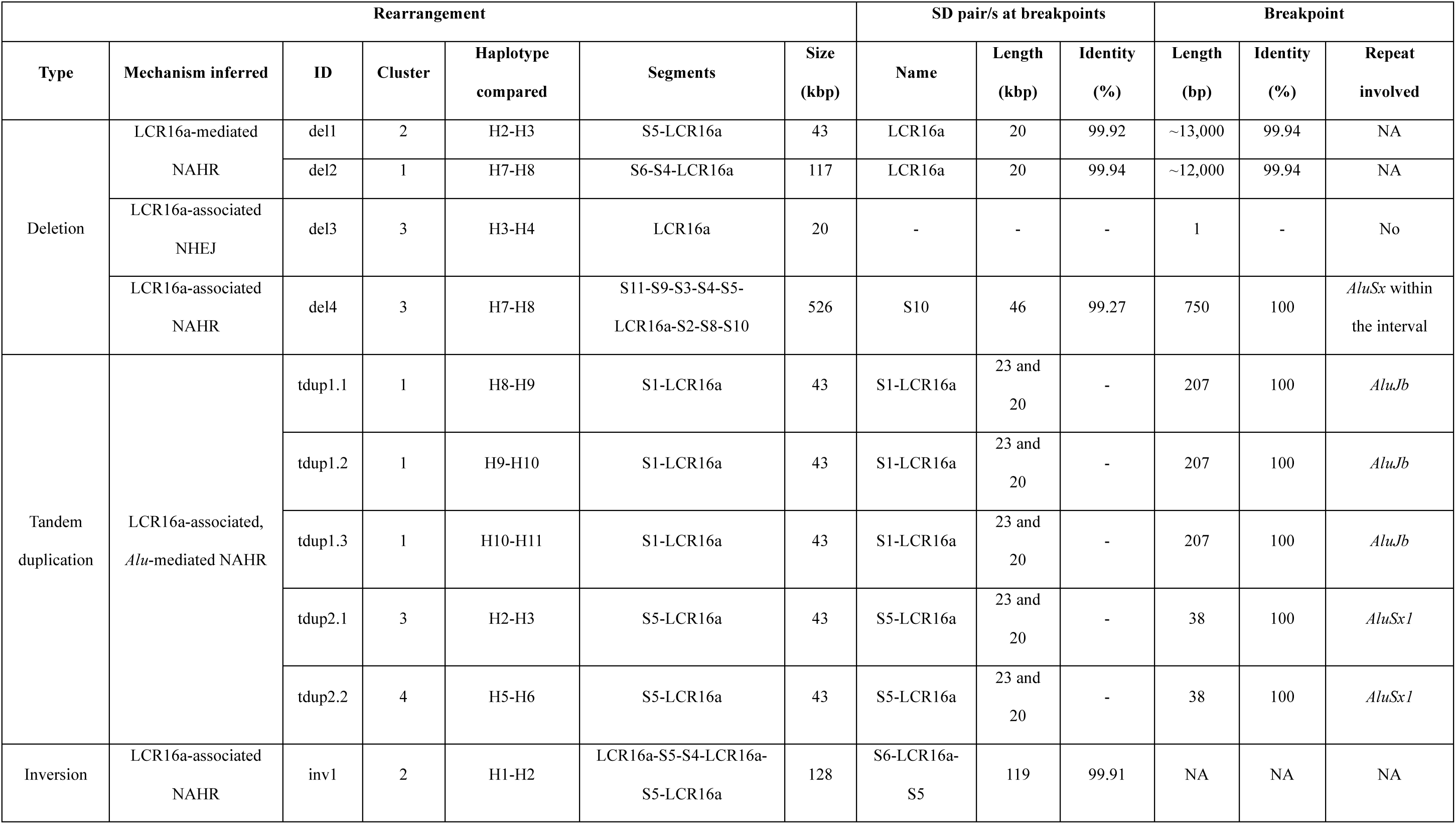
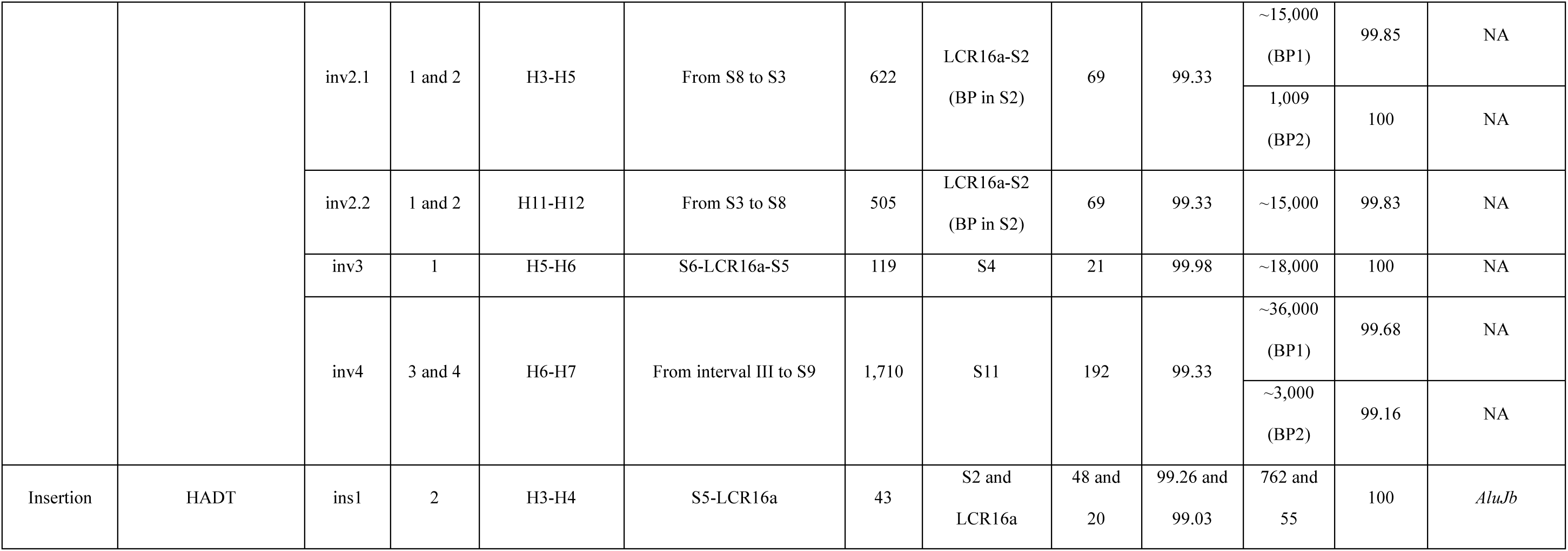
Pangenome 16p12.3–p13 structural variations and mechanisms of origin. BP: breakpoint; bp: base pairs.

We did not find the structure of the T2T-CHM13 reference (H8 haplotype) in any sequence from the pangenome dataset. The T2T-CHM13 configuration is simpler than H1, the most common haplotype for clusters 1-2 (12/39 analyzed sequences, 31%). In fact, the S6 segment (light blue in **Figure 2**) is present in two copies in most genomes (31/39, 79%) but is in single copy in H8.

In the subinterval spanning clusters 3 and 4, we identified eight alternative structures (H1–H8). In this case, the T2T-CHM13 reference has the most frequent H1 haplotype (17/35, 49%). Overall, structural variability primarily consisted in tandem duplications of segments S5-LCR16a in both clusters (tdup2), mirroring the tdup1 observed in cluster 1. Additionally, we identified two large structural variations: a 1.7 Mbp inversion in H7 (inv4) and a 526 kbp deletion in H8 (del4), each observed in a single genome.

We also classified a deletion of one LCR16a copy in H4 (del3) (**Figure 2**, **Table 1**).

### Majority of structural variation is generated by NAHR

We mapped breakpoint junctions of 15 different structural variations, including 10 copy number variations and five inversions (**Supplemental Data**). Breakpoint junction features revealed five distinct molecular mechanisms of structural variant formation (**Table 1**, **Figures 2, 3**). In seven cases, we could not refine the breakpoints that were assigned to kilobase-scale intervals within the duplicon/s potentially driving the rearrangement (**Table 1, Supplemental Data**).

Thirteen out of 15 events were driven by NAHR, with breakpoints located in three possible configurations: i) both breakpoints within a duplicon clustered with LCR16a (LCR16a-*associated* NAHR, 6/15 events); ii) both breakpoints within the LCR16a duplicon (LCR16a-*mediated* NAHR, 2/15 events); iii) one breakpoint within the LCR16a duplicon and the other within a LCR16a-adjacent duplicon (5/15 events). We assigned the third category to tdup1 and tdup2 events, where breakpoints were recurrent across haplotypes and copy-number expansions. These were refined to intervals of 207 and 38 bp, respectively, both corresponding to *Alu* sequences. Specifically, S1-LCR16a tandem duplications in cluster 1 (tdup1) were driven by NAHR between two *AluJb* elements, located at the start of S1 and at the end of LCR16a. Similarly, S5-LCR16a copy number variations in clusters 3 and 4 (tdup2) resulted from NAHR between two *AluSx1* elements at the start of S5 and at the end of LCR16a. We define this mechanism as LCR16a-associated, *Alu*-mediated NAHR (**Figures 2 and 3**, **Table 1**). The large deletion in clusters 3-4 (del4) and all five inversions (inv1, inv2.1, inv2.2, inv3, and inv4) were driven by LCR16a-associated NAHR, while deletions in clusters 1-2 (del1 and del2) result from LCR16a-mediated NAHR (**Figures 2 and 3**, **Table 1**).

**Figure 3.**
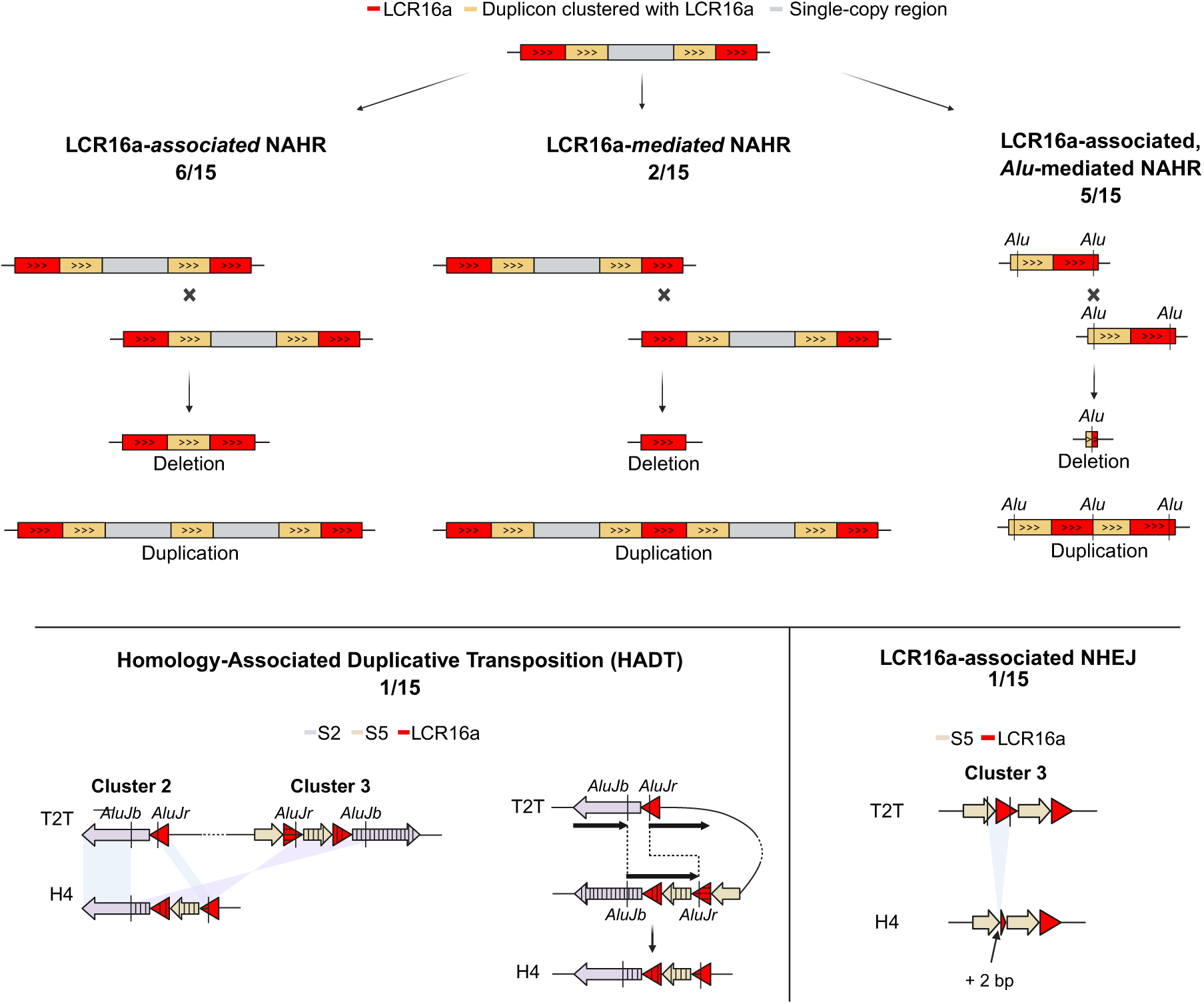
Models of mutational mechanisms in 16p12.3–p13.11 structural variation. (*top left*) LCR16a-associated NAHR involves recombination between paralogous duplicons clustered with LCR16a. (*top middle*) LCR16a-mediated NAHR involves recombination between LCR16a paralogous copies. (*top right*) LCR16a-associated, *Alu*-mediated NAHR involves crossing over at two *Alu* sequences, one located within the LCR16a duplicon and the other in a next to duplicon. This mechanism results in the deletion and duplication of the LCR16a and its associated duplicon. (*bottom left*) Homology-Associated Duplicative Transposition (HADT) involves a first recombination/switch at *AluJb* elements in inverted S1 copies and a second recombination/switch at *AluJr* elements in inverted LCR16a copies. The paralogous copies of the two inverted SD pairs are in clusters 2 and 3. During this process, a duplicative transposition of LCR16a and S5 occurs in cluster 2. (*bottom right*) LCR16a-associated NHEJ causes a deletion of S5 and LCR16a. The frequency of each mechanism is indicated. Created with BioRender.com.

Besides from these recurrent mechanisms, we identified a single rearrangement (del3, involving the deletion of one LCR16a copy accompanied by the insertion of two base pairs at the junction), whose breakpoint features are consistent with a non-homologous end joining (NHEJ) event (33) (**Figures 2 and 3**). We also identified a single insertion event in cluster 2 (ins1), whose breakpoint junction description is provided in the next section.

### *De novo*LCR16a-containing inverted duplicative transposition through iterative switching at *Alu* elements within two inverted SD pairs

We identified in the HG02145.2 genome a joint insertion of LCR16a and S5 segments (ins1 in cluster 2 of haplotype H4). To predict the mechanism underlying this duplicative transposition, we analysed the similarity between segments S2-LCR16a-S5-LCR16a at the end of H4 cluster 2 and all the homologous segments located in the four 16p12.3–p13.11 clusters from T2T-CHM13 reference. While the beginning of S2 and the end of the last LCR16a copy in H4 haplotype were more similar to their allelic copies, the central portion was more similar to the homologous segments mapped to cluster 3. These observations indicate that the insertion was generated by a complex mechanism involving two breakpoint junctions that correspond to four breakpoints in the parental sequence (**Figure 3**). Breakpoint junction mapping revealed two switches at *AluJ* elements located in two pairs of inverted SDs (S2 and LCR16a segments in clusters 2 and 3) that are ∼1 Mbp distant in the linear chromosome. We reason that, since the original sequence is still present in cluster 3 (unless there was a transposition from cluster 3 to 2 followed by a recombination event in the region between clusters 2 and 3 that restored the original cluster 3 sequence), a double recombination scenario can be ruled out as the origin of this insertion. Instead, the observed structure supports a replication-based mechanism involving two template switches occurring at homologous positions. We propose to define this mechanism as Homology-Associated Duplicative Transposition (HADT).

### Comparative analysis of 16p12.3–p13.11 architecture in primates

Given the recurrent involvement of LCR16a and *Alu* elements in the generation of 16p12.3–p13.11 genetic diversity among humans, we sought to investigate their contribution to shaping the structure of this region over longer timescales. To this aim, we compared the human T2T-CHM13 organization with the one at orthologous loci in other primates, by mapping the same intervals (single-copy regions I, II, and III; segments S1 to S11; LCR16a duplicon) to T2T genomes from the common chimpanzee (*Pan troglodytes*), Sumatran orangutan (*Pongo abelii*), and crab-eating macaque (*Macaca fascicularis*).

Comparison with the cow genome (bosTau9) suggests that the macaque sequence closely resembles the ancestral organization that existed ∼30 Mya. The macaque region is almost entirely devoid of SDs, except for a 17 kbp-long pair with 95% sequence identity. Both copies are located between segment S6 and single-copy interval II, oriented in the same direction, and separated by 17 kbp.

All segments, except S5—which uniquely maps to chr16:2 Mbp (16p13.3) orthologous region— uniquely map to the macaque 16p12.3–p13.11 orthologous region. Therefore, their positions in the macaque genome reflect their ancestral locations (**Figure 4**). In orangutan, the region has an extremely rearranged organization compared to the other primates, with a ∼4.5 Mbp long cluster 2, suggestive of orangutan lineage-specific structural changes.

**Figure 4.**
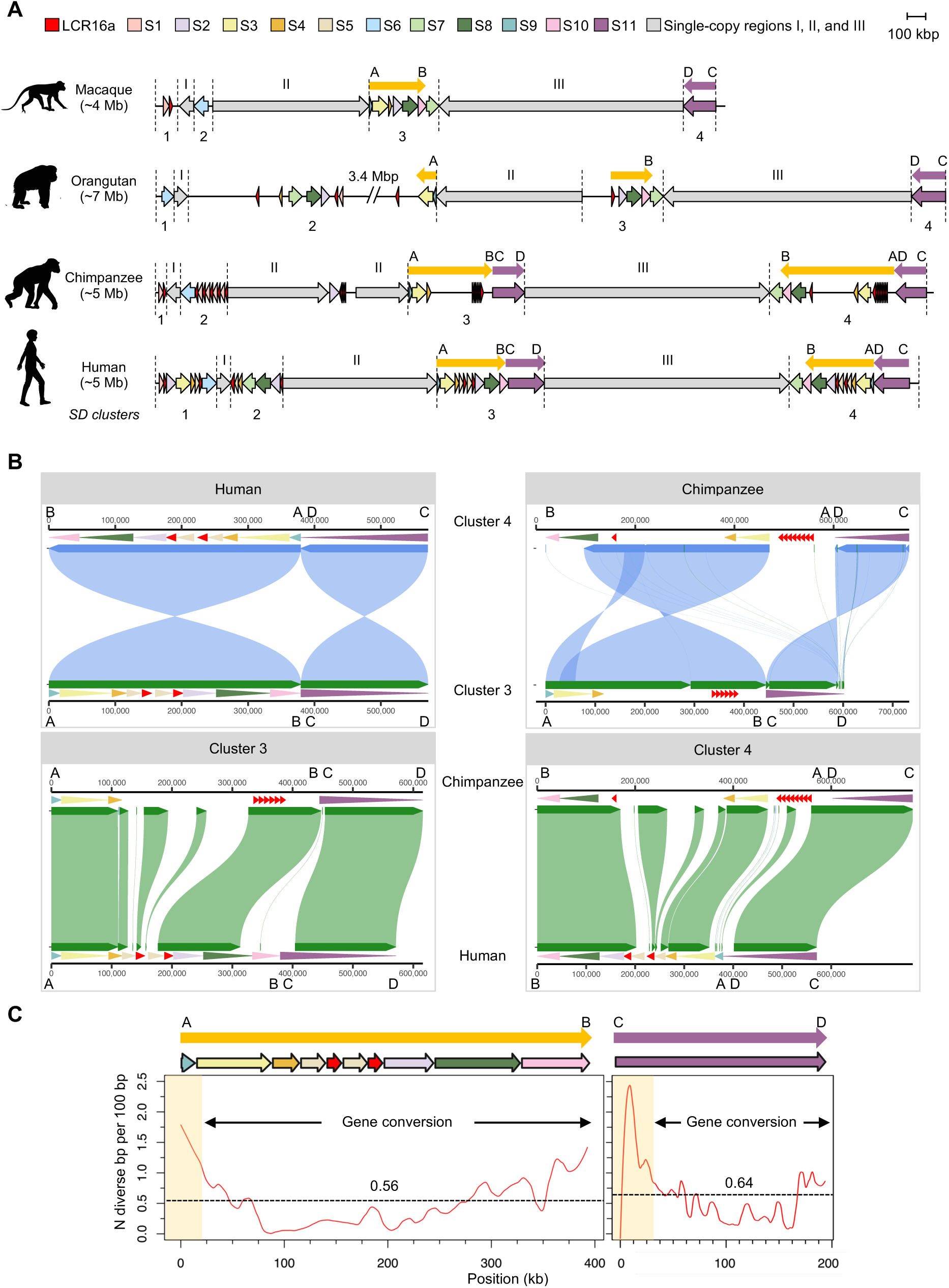
**16p12.3–p13.11 genomic architecture in primates**. A) Schematic of 16p12.3–p13.11 orthologous regions in macaque, orangutan, chimpanzee, and human. The size of the region is shown in parentheses. Vertical black dashed lines separate SD clusters from single-copy regions, respectively indicated by Arabic and roman numbers. The yellow and purple arrows highlight the two pairs of inverted segmental duplications shared by clusters 3 and 4 in human (AB and CD segments) and the positions of orthologous sequences in nonhuman primate genomes. B) (*top*) SVbyEye-visualization of the alignments between clusters 3 and 4 sequences of human (*left*) and chimpanzee (*right*). (*bottom*) SVbyEye-visualization of the alignments between human and chimpanzee cluster 3 and cluster 4 sequences, highlighting greater conservation in the outer regions and more divergence in the inner ones. C) Sliding-window diversity plot between human clusters 3 and 4 using sequences from 35 genomes. Horizontal dashed lines indicate the mean diversity across the interval. The portions not affected by interlocus gene conversion are highlighted in yellow.

In macaque and chimpanzee cluster 1, we identified only segments S1 and LCR16a, suggesting that the additional segments found in human cluster 1 arose through recent, human-specific duplicative transpositions. In agreement with previous studies (25, 26), the only LCR16a copy present in the macaque genome is mapped at cluster 1, next to the S1 segment. Conversely, in the chimpanzee and orangutan genomes we mapped, respectively, 25 and four LCR16a copies across the entire 16p12.3– p13.11 orthologous regions.

While macaque cluster 2 is composed of only one S6 copy, in chimpanzee we identified an expansion by tandem duplication of the S1-LCR16a segments (**Figure 4A**). We observe that in the majority of human haplotypes (24 out of 39, 61%), interval I and segment S6 are in the same orientation as in macaque and chimpanzee genomes (with S6 belonging to cluster 2). This arrangement reflects the ancestral organization and is inverted compared to the orientation in T2T-CHM13 and orangutan genomes (**Figures 2 and 4A**). Similarly to cluster 1, human cluster 2 was the acceptor region of recent, human-specific duplicative transpositions.

As previously specified, human clusters 3 and 4 contain the same segments but in a different configuration (**Figures 1 and 4**). These segments, except part of S4 and S5, are already present at the macaque orthologous region, however segments from S9 to S10 (“AB”, yellow arrow) are only present in macaque cluster 3 orthologous region, while segment S11 (“CD”, purple arrow) is only mapped to macaque cluster 4 orthologous region. We observe a similar configuration in orangutan with only S11 segment mapped to cluster 4, although orangutan-specific rearrangements, including an inversion, modified the organization of the region from cluster 1 to cluster 3. Moreover, the single-copy region III (the one between clusters 3 and 4) is inverted in human and chimpanzee compared to macaque and orangutan, with the latter therefore corresponding to the ancestral configuration.

Chimpanzee organization of clusters 3 and 4 also mirrors the human structure, with sequence blocks switched in position and inverted between the two clusters (AB-CD/BA-DC) (**Figure 4B**, top). While the extremities of each cluster are conserved between human and chimpanzee, there is a higher interspecies divergence in the central portions (**Figure 4B**, bottom). Overall, this suggests that the event/s generating clusters 3 and 4 likely occurred in the human-chimpanzee common ancestor. After the split, in each lineage clusters 3 and 4 were further modified and expanded through interspersed and tandem duplications. This instability was higher in the chimpanzee lineage, where the clusters experienced, among other events, tandem duplications of LCR16a (**Figure 4A, B**).

Human SDs in clusters 3 and 4 are highly similar (identity of 99.33% for the “AB” yellow pair; identity of 99.46% for the “CD” purple pair), mirroring the high similarity between chimpanzee clusters 3 and 4 SDs (99.20 and 99.65% identity for the “AB” and “CD” pairs, respectively). When we measured the diversity along the “AB” yellow and “CD” purple intervals between human clusters 3 and 4 sequences, we found that the level of diversity is similar between the two intervals and central portions of both intervals show a lower diversity compared to the extremities, because of interlocus gene conversion (**Figure 4C**). This pattern is the opposite of what is observed in the interspecies comparison of orthologous copies (**Figure 4B**).

### Dating of interspersed duplications shaping human 16p12.3–p13.11 SD clusters

To further elucidate the events shaping the complex architecture of the 16p12.3–p13.11 locus in humans, we estimated the timing of interspersed duplications that converted an ancestral SD-free region into its current SD-rich state. As 16p12.3–p13.11 paralogous segments have been extensively homogenized by interlocus gene conversion (34), we selected intervals not affected by this process in any human haplotype that would otherwise cause an underestimation of the duplication timings.

We estimated that S7 segment duplicated from cluster 4 to cluster 2 around 3.6–3.9 Mya and S3 duplicated from cluster 3 to cluster 1 around 2.2–2.3 Mya (**Supplemental dataset**). When we considered the two large duplications shared by clusters 3 and 4 (yellow and purple arrows in **Figure 1**), we could use only two intervals: a 20.5 kbp region at the tail of the “yellow” arrow and a 27 kbp region at the tail of the “purple” arrow, as these are the only portions not affected by interlocus gene conversion (**Figure 4C**). These duplications were dated to 6.0–6.5 and 5.9–6.3 Mya, respectively, suggesting that they likely occurred simultaneously in the human-chimpanzee common ancestor shortly before their split (**Supplemental dataset**).

### Model of 16p12.3–p13.11 structural evolution

We integrated data from primate genome alignments, duplication timing estimations, and breakpoint mapping to build a model of the evolutionary structural changes that brought to the current 16p12.3– p13.11 architecture in humans. We hypothesize that 30 Mya the locus had a structure similar to that observed in the macaque genome. By comparing human and macaque cluster 3 sequences, we found that a 9.5 kbp segment located between S4 and S2 segments and flanked by an *AluSx1* and a *L1PA7* LINE element was deleted and replaced by an LCR16a copy. As human, chimpanzee, and orangutan have at least one LCR16a copy at this position, we presume that this event occurred in the great ape common ancestor ∼20 Mya (**Figure 5**). In the human-chimpanzee common ancestor, a 38 kbp duplicative transposition expanded cluster 3 with sequences (corresponding to part of S4 and the entire S5 segment) derived from the 16p13.3 (chr16:2Mbp) region (**Figure 5**).

**Figure 5.**
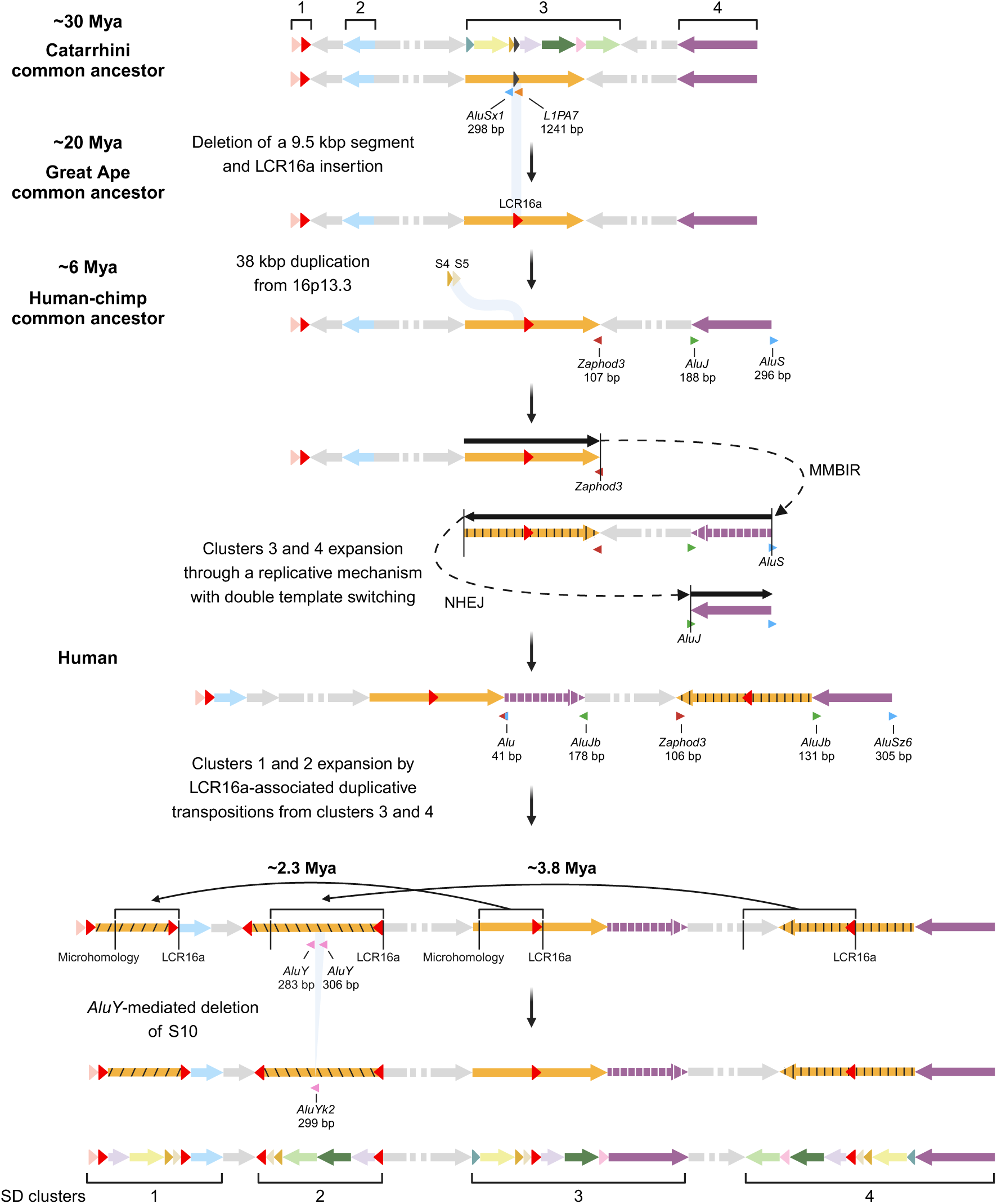
Model of 16p12.3–p13.11 structural evolution. Structural changes that shaped the human 16p12.3– p13.11 from the organization inferred in the Catarrhini common ancestor (∼30 Mya) to the T2T-CHM13 structure. The model was reconstructed based on segment mapping to primate T2T genomes, phylogenetic analysis, and breakpoint examination. In the Catarrhini common ancestor, the region was devoid of SDs. A detailed structure with all segments and a simplified structure with a single yellow arrow including all modules of cluster 3 (except S7 green segment) are shown. In the great ape common ancestor, a 9.5 kbp segment in cluster 3 was deleted and replaced by an LCR16a copy. In the human-chimpanzee common ancestor, there was a 38 kbp duplication from 16p13.3 (chr16:2Mb) to cluster 3 and a double template switching event that expanded clusters 3 and 4 together with the inversion of the region in-between. In the last 4 million years of human evolution, clusters 1 and 2 expanded through at least two LCR16a-associated duplicative transpositions from clusters 3 and 4. Recombination between two *AluY* elements led to a 45 kbp deletion in cluster 2. Created with BioRender.com.

Sequence analysis of the breakpoint junction between the yellow and purple arrows in human cluster 3 revealed a 41 bp incomplete *Alu* element. This sequence originates from ancestral full-length *Zaphod3* and *AluS* elements, which are respectively located at the head of the yellow arrow and the tail of the purple arrow in cluster 4 (**Figure 5**). We observed a 3 bp microhomology (AAG) between the *Zaphod3* and *AluS* elements at the breakpoint junction. Similarly, we identified a 131 bp truncated *AluJb* at the breakpoint junction between the yellow and purple arrows in human cluster 4, which derives from the ancestral *AluJ* sequence at the head of the purple arrow in human cluster 3 (**Figure 5**). At this breakpoint junction, we observed no homology between the sequences.

Based on our time estimates and breakpoint junction analysis, we reasoned that clusters 3 and 4 were formed by a single mutational event characterized by two template switches during DNA replication, in the human-chimpanzee common ancestor ∼6 Mya (**Figure 5**). We hypothesize that a first template switch occurred at the *Zaphod3* element located at the head of the yellow arrow. Consequently, DNA replication was restored at the *AluS* sequence at the tail of the purple arrow and the purple block, the single-copy segment III, and the yellow block were copied in the opposite direction. A second template switch restored the original replication direction. This second time, the fork resumed replication at an *AluJ* element located at the head of the purple arrow (**Figure 5**). This complex rearrangement generated at the same time the duplications in clusters 3 and 4, where the boundaries between the yellow and purple segments are characterized by incomplete *Alu* elements, and the inversion of the single-copy region in-between. The first template switch was likely mediated by an MMBIR (microhomology-mediated break induced replication) mechanism, while the second occurred via NHEJ, involving a dinucleotide (TC) insertion before the truncated *AluJ* element.

Clusters 1 and 2 expanded more recently in the human lineage by means of at least two duplicative transpositions. The first one occurred around 3.8 Mya from cluster 4 to cluster 2 and was featured by the presence of an LCR16a copy at one extremity and no homology with the target site at the other. A subsequent duplicative transposition from cluster 3 to cluster 1, around 2.3 Mya, was featured by the presence of an LCR16a copy at one extremity and microhomology (CCT) with the target site at the other. Finally, we inferred a 45 kbp deletion of S10 segment in cluster 2 driven by *AluY*-mediated NAHR (**Figure 5**).

## DISCUSSION

Human and great ape genomes are characterized by interspersed SDs (16), as opposed to the SD tandem pattern observed in other mammalian genomes (35). While the propagation of tandem SDs can be explained by NAHR, the spread of interspersed SDs does not have an immediate mechanistic explanation. SD expansion in the human genome has been associated with the presence of *Alu* elements and core duplicons (17, 21). However, the specific role of these sequences in the mechanisms of SD generation remains unresolved.

In this study, we addressed this question by analysing intraspecies and evolutionary structural variation at the 16p12.3–p13.11 locus. This region contains the ancestral copy of the LCR16a core duplicon, a genomic segment central to the formation of the SD architecture on the short arm of chromosome 16. This locus is rich in intrachromosomal and interspersed SDs and is medically relevant as it is the site of recurrent pathogenic copy-number variations mediated by highly identical SDs. We catalogued structural rearrangements within this interval, mapped their breakpoints, and inferred their underlying mechanisms of origin.

Our analysis revealed that 16p12.3–p13.11 SD-mediated instability occurs through multiple recombination/replication-based mechanisms, with the LCR16a core duplicon involved in the majority of processes, and *Alu* elements often playing a key role. The majority of intraspecies SD expansion arises by NAHR, generating novel copies in a tandem configuration. We note that while NAHR with breakpoints within LCR16a-associated duplicons has generated all five inversions, NAHR with breakpoints within LCR16a itself causes copy-number variations. This instability increases when the breakpoints occur at *Alu* elements within LCR16a and an adjacent duplicon (LCR16a-associated, *Alu*-mediated NAHR), generating multiple and polymorphic tandem copies of the LCR16a-duplicon pair, as observed for segments S1-LCR16a in cluster 1 and S5-LCR16a in clusters 3 and 4.

The structural variation of cluster 2 in haplotype H4 was the only case showing an interspersed duplication, with the joint insertion of duplicons LCR16a and S5, copied from sequences ∼1 Mbp away in cluster 3. This structure offered the unique possibility to witness and gain insights into the formation of novel interspersed, inverted SD pairs that include the LCR16a duplicon. Two breakpoint junctions were generated during this rearrangement and correspond to *AluJ* elements within two pairs of inverted SDs (LCR16a and S2). We suppose that this duplicative transposition was generated through a replicative mechanism with double template switching, where *AluJ* sequences within two pairs of homologous inverted SDs acted as substrates for recombination (HADT mechanism). A double template switching mechanism has been proposed to explain the origin of the duplication-triplication/inverted-duplication complex rearrangement (36). These cases involved the first switch at a pair of inverted intrachromosomal SDs, while the second switch was at non-homologous positions, differently from the H4 case where both switches occurred at pairs of inverted SDs.

We identified a similar mechanism forming at the same time two pairs of large, inverted duplications at clusters 3 and 4, together with the inversion of the intervening sequence, in the human-chimpanzee common ancestor. This is an example of a drastic and rapid evolutionary change in the genome, driven by a single complex rearrangement that created novel pairs of inverted SDs and a large inversion. We propose that this rearrangement derived from two template switches, the former at tracts with microhomology—reminding the fork stalling template switching (FoSTeS)/MMBIR mechanisms (37, 38)—and the latter at nonhomologous positions, suggestive of NHEJ (**Figure 5**). Notably, in both switches the fork resumes replication at an *Alu* element (*AluS* and *AluJ* at the purple segment edges). Several lines of evidence support this model: i) the similar dating of the two pairs of inverted duplications (yellow AB and purple CD segments); ii) the presence of only two breakpoint junctions; and iii) the inverted orientation of the sequence in-between in the majority of humans and in chimpanzee sequence, compared to orangutan and macaque structures. Because of the conformation of the two pairs of inverted duplications (AB/CD in human cluster 3 and BA/DC in human cluster 4), a previous study suggested a concomitant origin for these duplications through a circular DNA intermediate (29). However, this mechanism would require two steps: first, the inversion of the region spanning S7-interval III-S11; second, the circularization of the DNA from S9 to S11, followed by its reinsertion at the tail of the S7 segment in cluster 4, precisely at one of the two breakpoints of the previous inversion. While possible, this scenario is less parsimonious. Another alternative involves two duplications (yellow segment from cluster 3 to cluster 4 and purple segment from cluster 4 to cluster 3), along with the inversion of S7-interval III. However, this too is less likely. Both of these multi-step scenarios would require the exact reuse of the same breakpoints, making them less probable.

Although the two SDs pairs in clusters 3 and 4 originated in the human-chimpanzee common ancestor, extensive interlocus gene conversion has homogenized the sequences within each species, while increasing interspecies differences of orthologous copies, a pattern recently reported for SDs at the genome-wide level (34). This pattern is more pronounced in the middle of both sequence pairs, consistent with a higher frequency of interlocus gene conversion toward the inner regions of duplicate sequences (39) (**Figure 4B, C**).

The possible replicative mechanisms driving H4 duplicative transposition and cluster 3-4 formation align with previous findings in humans and budding yeast, reporting that duplications arise from replication stress and repair of broken forks via homology-dependent and-independent replication-based processes (40, 41). While it remains unclear if this is the primary mechanism of interspersed SD formation in human and primate genomes, it may explain other focal SD accumulations across the human genome. Although less frequent than NAHR, replicative mechanisms have a greater impact on subsequent genome instability, as they produce interspersed and inverted SDs. These, in turn, can trigger inversions and copy-number changes through NAHR, affecting large intervals that may contain dosage sensitive genes, ultimately leading to disease (42).

This study highlights the synergistic interplay between *Alu* elements and LCR16a as key drivers of genomic instability and SD expansion at the 16p12.3–p13.11 locus. Structural rearrangements arise through both replication-and recombination-based mechanisms, with *Alu* elements acting as preferential sites for strand exchange. This instability creates structural diversity among humans and has shaped the dynamic architecture of 16p12.3–p13.11 over the past 30 million years, making it susceptible to pathogenic rearrangements. Future research will explore whether similar mechanisms operate in regions containing other core duplicons—some of the most dynamic and unstable parts of the human genome. These regions have provided abundant raw material for gene innovation in recent human evolution, while creating predispositions to neurodevelopmental diseases (43).

## METHODS

### Structural variation analysis

We analyzed structural variation using an approach similar to the one previously applied to the 10q11.22 chromosomal region (22). We obtained SD annotation from the relative track in the UCSC (University California Santa Cruz) genome browser. We searched for intervals corresponding to the 16p12.3–p13.11 chromosomal region in diploid and phased assemblies of 47 samples produced in the frame of the Human Pangenome Project (32), using minimap2 v2.28 (default parameters) (44) and reference sequences from T2T-CHM13v2.0 as query. Human pangenome data analysed during this study are publicly available from https://github.com/human-pangenomics/HPP_Year1_Data_Freeze_v1.0.

Figure 2 was obtained in R v4.3.2 (45) using the ggplot2 v3.5.1 (46) and gggenomes v1.0.1 (47) packages.

To define breakpoint regions, we identified the SD pair/s that putatively mediated the structural variant formation. We multialigned the SD sequence/s from the haplotype under investigation with those from the putative parental haplotype using MAFFT (48). We calculated pairwise nucleotide diversity in 500 bp sliding windows with a 100 bp increment by using PopGenome R package v2.7.7 (49). We visually identified the breakpoint as the position where the derived hybrid SDs become more similar from one copy to the other of the same pair from the parental haplotype. We mapped the identified breakpoint regions in the T2T-CHM13 (v2.0) genome by BLAT (50) and assessed repetitive element content (RepeatMasker Repetitive Elements UCSC track). Figure 3 was created with BioRender (https://www.biorender.com).

### Evolutionary analysis

We downloaded chimpanzee (mPanTro3) and orangutan (mPonAbe1) T2T genomes from https://www.genomeark.org/t2t-all and macaque T2T genome (T2T-MFA8) from https://github.com/zhang-shilong/T2T-MFA8 (51). We aligned the entire 16p12.3–p13.11 human sequence and the sequences of the various segments to the primate genomes using minimap2 v2.28 (default parameters). We used ggplot2 v3.5.1 and gggenomes v1.0.1 packages to create Figure 4A.

In the phylogenetic analysis, we assessed segment intervals not affected by interlocus gene conversion (52). We used a 13 kbp interval to analyze S7 duplications and a 4 kbp interval for S2 duplications. We multialigned human, orangutan, and macaque sequences with MAFFT. We built phylogenetic trees by using the Maximum Likelihood method and Kimura 2-parameter model (53) with the complete deletion option, in MEGA11 (54). Statistical significance of nodes was evaluated by performing the bootstrap test with 100 replicates (55). To estimate duplication timing, we considered macaque as outgroup and the human-orangutan split at 18.2–19.6 Mya (16).

We aligned human and chimpanzee cluster 3 and 4 sequences with minimap2 v2.28 (-x asm20-c –eqx parameters) and visualized the alignment using ggplot2 v3.5.1 and SVbyEye v0.99.0 (56) R packages. We calculated the diversity between clusters 3 and 4 sequences from 35 human genomes in 100 bp sliding windows with a 10 bp increment by using the PopGenome R package v2.7.7.

## ACKNOWLEDGMENTS

This study received funding from the European Union – Next Generation EU – National Recovery and Resilience Plan (NRRP), Mission 4, Component 2, Investment 1.1, call PRIN 2022 D.D. 104 02.02.2022 to G.G. (2022RB88C7).

## Authors’ contributions

G.G. conceived and supervised the study. S.F. and G.G. analysed and interpreted the data. S.F. and G.G. wrote the manuscript.

## ADDITIONAL FILES

**Supplemental Table 1**

**Supplemental Data**

**Supplemental Dataset**

