## Supplemental Data for "Synergistic role of *Alu* and core duplicon sequences in driving genomic instability at the disease-associated 16p12.3–p13.11 region"

#### Contents

1. H1-H2 inv1 (cluster 2) breakpoint analysis (Supplemental Figure S1) …..……..… Pag. 2

2. H3-H5 inv2.1 (clusters 1-2) breakpoints analysis (Supplemental Figure S2) …..…. Pag. 3

3. H5-H6 inv3 (cluster 1) breakpoints analysis (Supplemental Figure S3) ………..…. Pag. 4

4. H6-H7 inv4 (clusters 3-4) breakpoints analysis (Supplemental Figure S4) ……….. Pag. 5

5. H2-H3 del1 (cluster 2) breakpoint analysis (Supplemental Figure S5) …………..... Pag. 6

6. H7-H8 del2 (cluster 1) breakpoint analysis (Supplemental Figure S6) …………..... Pag. 7

7. H7-H8 del4 (cluster 3) breakpoint analysis (Supplemental Figure S7) …………..... Pag. 8

8. Tandem duplications (cluster 1) breakpoints analysis (Supplemental Figure S8) …. Pag. 9

9. Tandem duplications (clusters 3-4) breakpoint analysis (Supplemental Figure S9) .. Pag. 11

### 1. H1-H2 inv1 (cluster 2) breakpoint analysis

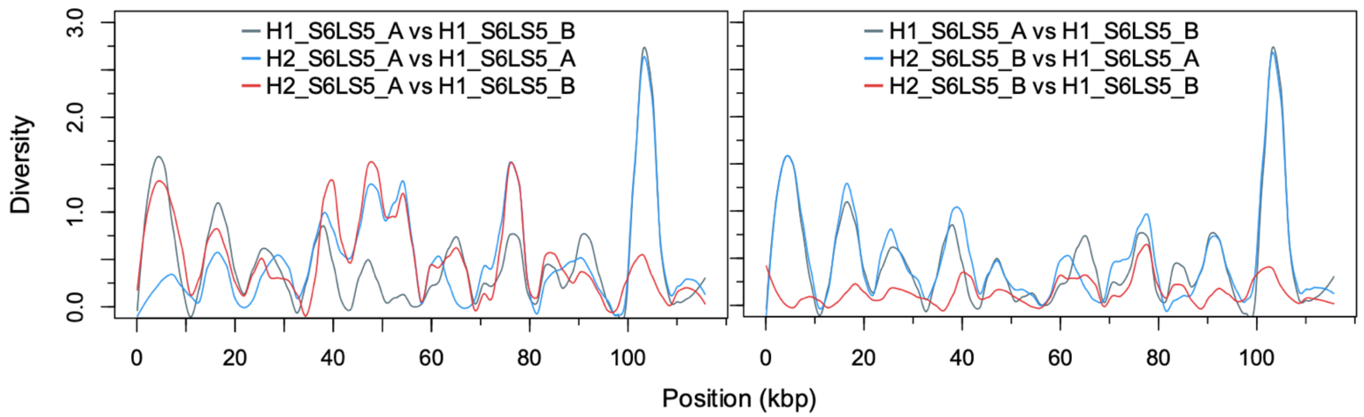


**Supplemental Figure S1.**

Diversity plots of H2 (HG02717.1) breakpoint regions compared with H1 (HG002.1) S6, LCR16a (L), and S5 putative SDs mediating the inversion inv1 in cluster 2. The plots show pairwise nucleotide diversity in 500 bp sliding windows with 100 bp increments. In the left graph, the blue and red lines represent comparisons of the H2 first breakpoint region (H2_S6LS5_A) with H1 parental SDs (H1_S6LS5_A and H1_S6LS5_B, respectively). In the right graph, the blue and red lines represent comparisons between the H2 second breakpoint region (H2_S6LS5_B) with H1 parental SDs (H1_S6LS5_A and H1_ S6LS5_B, respectively). The gray line serves as a reference, comparing H1 parental SDs (H1_S6LS5_A vs H1_ S6LS5_B).

The exact breakpoint could not be determined.

### 2. H3-H5 inv2.1 (clusters 1-2) breakpoints analysis

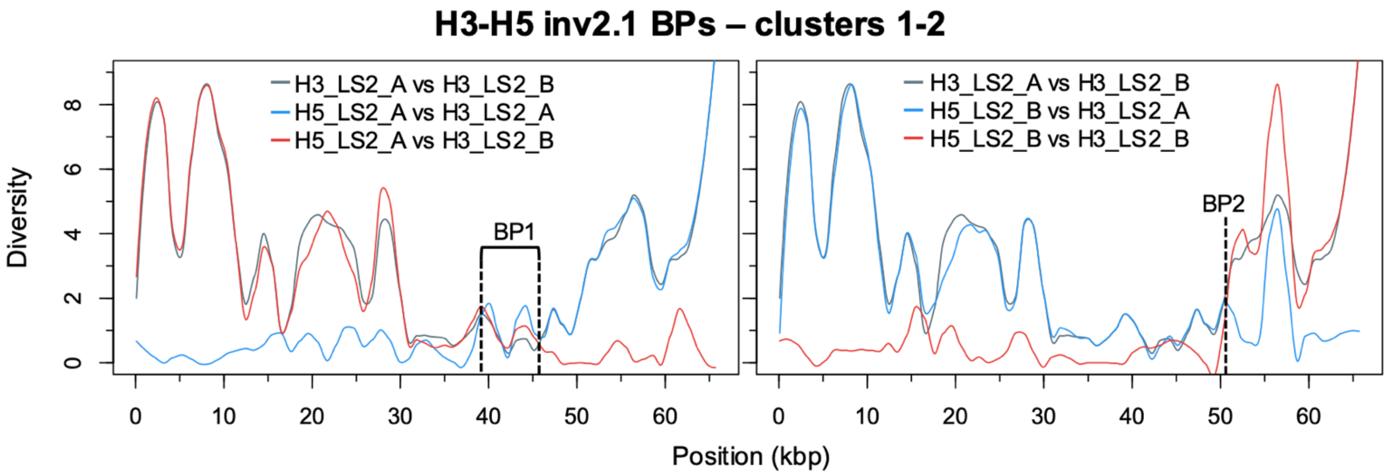


**Supplemental Figure S2.**

Diversity plots of H5 (HG01891.1) breakpoint regions compared with H3 (HG02886.1) LCR16a (L) and S2 putative SDs mediating the inversion inv2.1 in clusters 1-2. The plots show pairwise nucleotide diversity in 500 bp sliding windows with 100 bp increments. In the left graph, the blue and red lines represent comparisons of the H5 first breakpoint region (H5_LS2_A) with H3 parental SDs (H3_LS2_A and H3_LS2_B, respectively). In the right graph, the blue and red lines represent comparisons of the H5 second breakpoint region (H5_LS2_B) with H3 parental SDs (H3_LS2_A and H3_LS2_B, respectively). The gray line serves as a reference, comparing H3 parental SDs (H3_LS2_A and H3_LS2_B).

In this case, we identified switches at different positions within the two breakpoint regions. The final breakpoint regions are indicated as BP1 (~15 kbp) and BP2 (1,009 bp).

#### 3. H5-H6 inv3 (cluster 1) breakpoints analysis

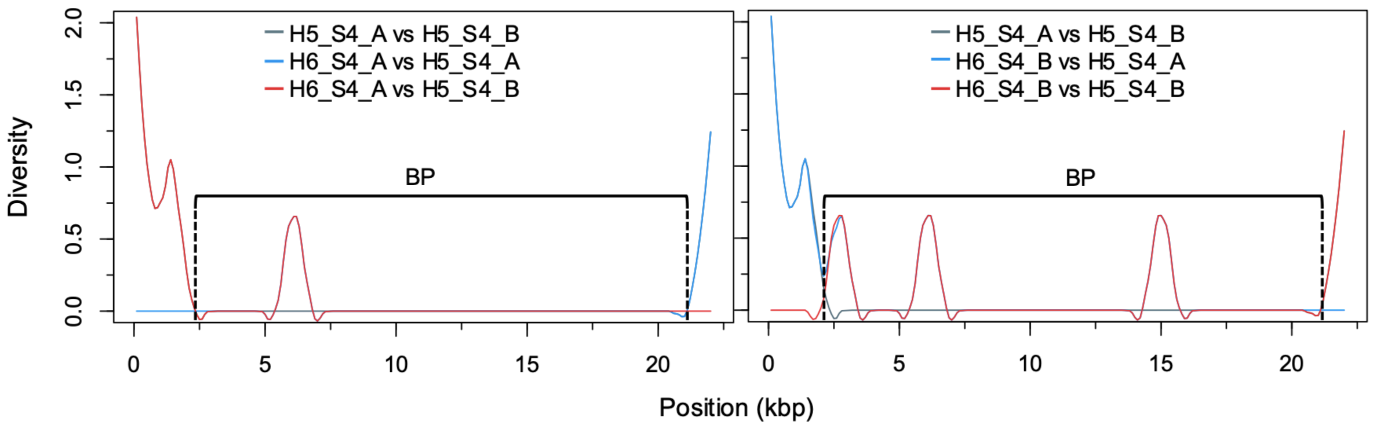


**Supplemental Figure S3.**

Diversity plots of H6 (HG02559.1) breakpoint regions compared with H5 (HG01891.1) S4 putative SDs mediating the inversion inv3 in cluster 1. The plots show pairwise nucleotide diversity in 500 bp sliding windows with 100 bp increments. In the left graph, the blue and red lines represent comparisons of the H6 first breakpoint region (H6_S4_A) with H5 parental SDs (H5_S4_A and H5_S4_B, respectively). In the right graph, the blue and red lines represent comparisons of the H6 second breakpoint region (H6_S4_B) with H5 parental SDs (H5_S4_A and H5_S4_B, respectively). The gray line serves as a reference, comparing H5 parental SDs (H5_S4_A and H5_S4_B).

The final breakpoint region was consistent across both plots and corresponds to the majority of S4 SDs (~18 kbp).

4. **H6-H7 inv4 (clusters 3-4) breakpoints analysis**

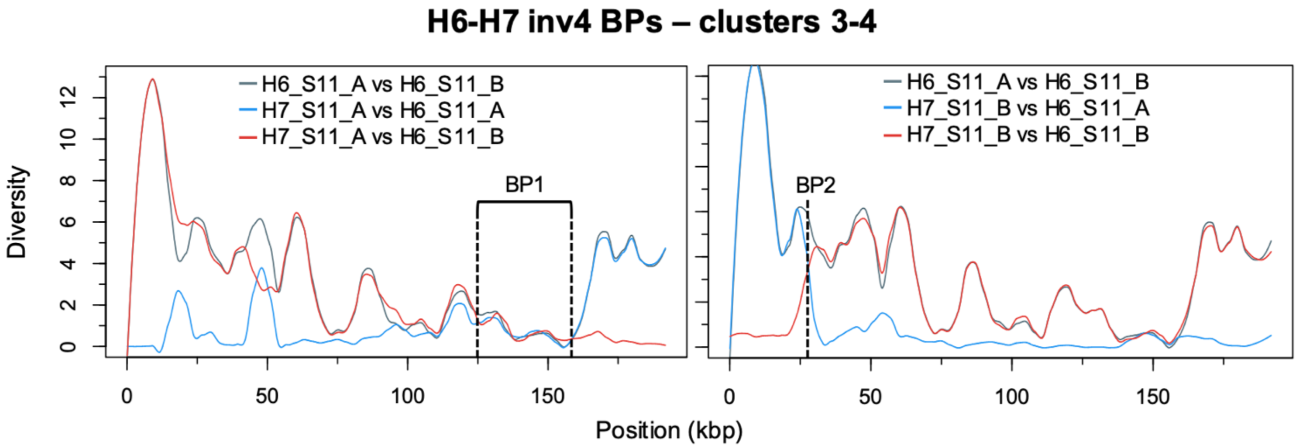


**Supplemental Figure S4.**

Diversity plots of H7 (HG01891.2) breakpoint regions compared with H6 (HG02886.2) S11 putative SDs mediating the inversion inv4 in clusters 3-4. In the left graph, the blue and red lines represent comparisons of the H7 first breakpoint region (H7_S11_A) with H6 parental SDs (H6_S11_A and H6_S11_B, respectively). In the right graph, the blue and red lines represent comparisons of the H7 second breakpoint region (H7_S11_B) with H6 parental SDs (H6_S11_A and H6_S11_B, respectively). The gray line serves as a reference, comparing H6 parental SDs (H6_S11_A and H6_S11_B).

In this case, we identified switches at different positions within the two breakpoint regions. The final breakpoint regions are indicated as BP1 (~36 kbp) and BP2 (~3 kbp).

### 5. H2-H3 del1 (cluster 2) breakpoint analysis

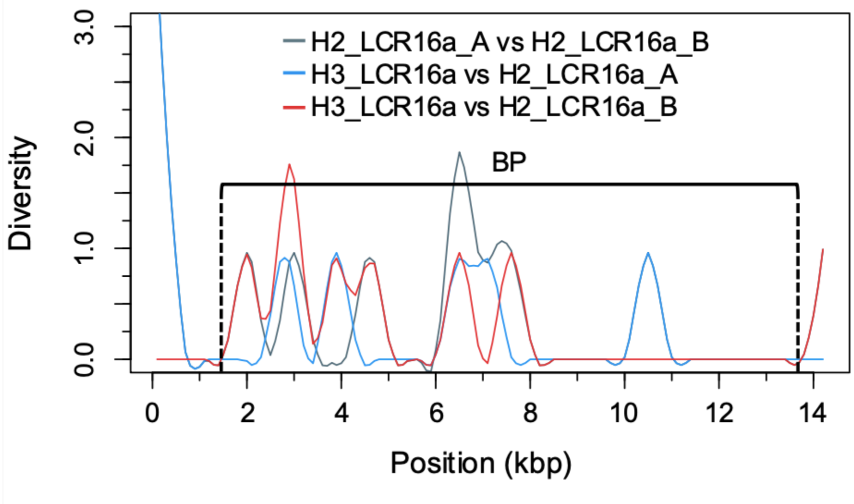


**Supplemental Figure S5**

Diversity plots of H3 (HG02886.1) breakpoint region compared with H2 (HG02717.1) LCR16a putative SDs mediating the deletion del1 in cluster 2. The plot shows pairwise nucleotide diversity in 500 bp sliding windows with 100 bp increments. The blue and red lines represent comparisons of the H3 breakpoint region (H3_LCR16a) with H2 parental SDs (H2_LCR16a_A and H2_LCR16a_B, respectively). The gray line serves as a reference, comparing H2 parental SDs (H2_LCR16a_A and H2_LCR16a_B).

The final breakpoint region corresponds to the majority of LCR16a SDs (~13 kbp).

6. **H7-H8 del2 (cluster 1) breakpoint analysis**

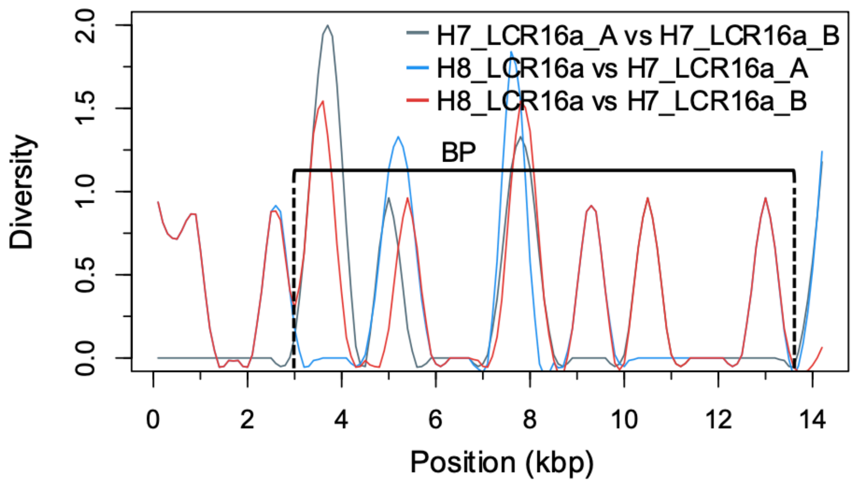


**Supplemental Figure S6.**

Diversity plots of H8 (T2T-CHM13) breakpoint region compared with H7 (HG03486.2) LCR16a putative SDs mediating the deletion del2 in cluster 1. The plot shows pairwise nucleotide diversity in 500 bp sliding windows with 100 bp increments. The blue and red lines represent comparisons of the H8 breakpoint region (H8_LCR16a) with H7 parental SDs (H7_LCR16a_A and H7_LCR16a_B, respectively). The gray line serves as a reference, comparing H7 parental SDs (H7_LCR16a_A and H7_LCR16a_B).

The final breakpoint region corresponds to the majority of LCR16a SDs (~12 kbp).

### 7. H7-H8 del4 (cluster 3) breakpoint analysis

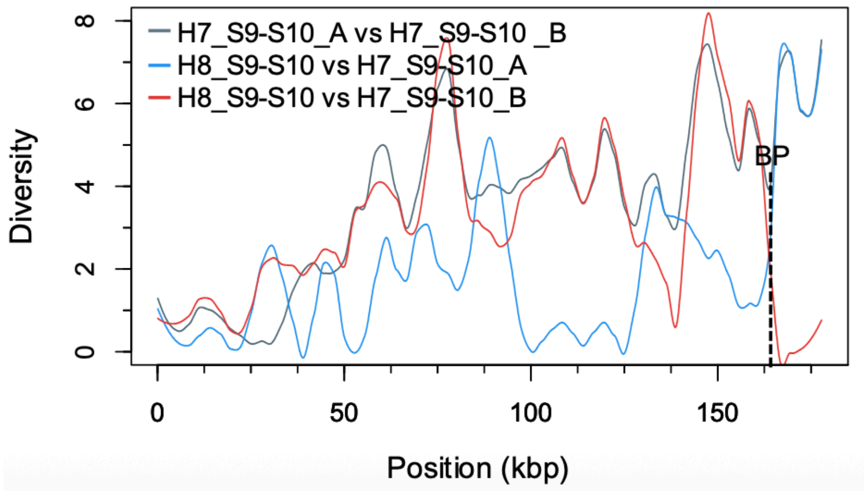


**Supplemental Figure S7.**

Diversity plot of H8 (HG01243.1) breakpoint region compared with H7 (HG01891.2) from S9 to S10 putative SDs mediating the deletion del4 in cluster 3. The plot shows pairwise nucleotide diversity in 500 bp sliding windows with 100 bp increments. The blue and red lines represent comparisons of the H8 breakpoint region (H8_S9-S10) with H7 parental SDs (H7_S9-S10_A and H7_S9-S10_B, respectively). The gray line serves as a reference, comparing H7 parental SDs (H7_S9-S10_A and H7_S9-S10_B).

The final breakpoint (750 bp) is represented by the dotted line.

### 8. Tandem duplications (cluster 1) breakpoints analysis

A

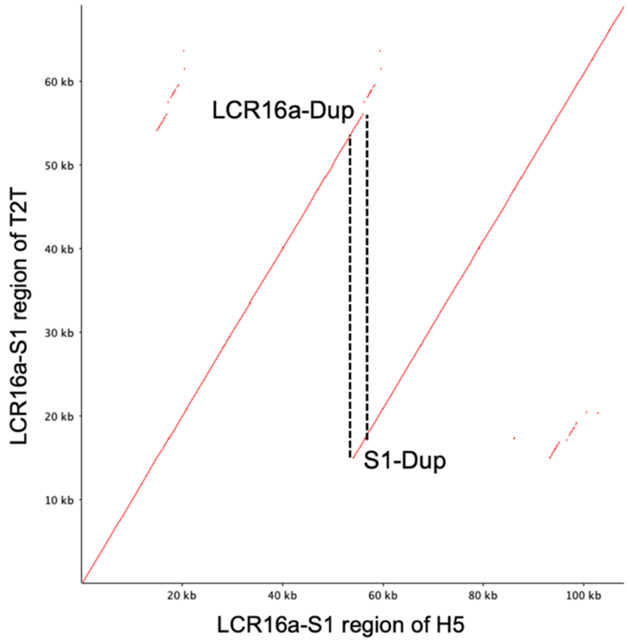


B

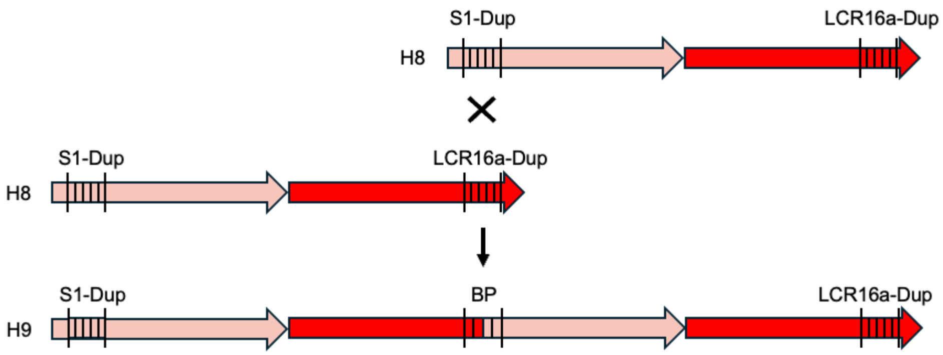


C

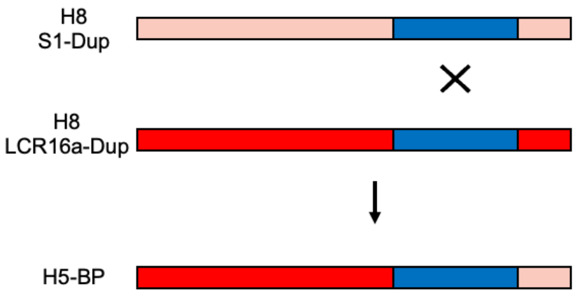


**Supplemental figure S8.**

To identify the breakpoints of the tandem duplications in cluster 1, we used a dot-plot comparison between the S1-LCR16a segments of H8 (T2T) reference genome and the region containing all multiple copies of S1-LCR16a from the haplotypes where these duplications occur.

An example of a dot-plot comparing H8 S1-LCR16a and H9 S1-LCR16a-S1-LCRC16 is shown in the figure. We observed that the start of S1 contains a sequence, named as S1-Dup, which is identical to a sequence found at the end of LCR16a, named as LCR16a-Dup (panels A and B in Supplemental figure S8). In H9, this sequence was present in three copies: one at the start of S1, one at the end of LCR16a, and one in the region between the first LCR16a and the second S1, which was named BP. This pattern is consistent with duplications mediated by NAHR, where the mediating segmental duplication is typically present in three copies within the rearranged haplotype. The same pattern was observed across all haplotypes containing these tandem duplications.

To further investigate, we retrieved the sequences corresponding to S1-Dup, LCR16a-Dup, and BP from all the rearranged haplotypes, aligned them using MAFFT (Katoh et al. 2009), and manually inspected the alignments.

We found that the first part of H9-BP is more similar to H8 LCR16a-Dup, while the final part is more similar to H8 S1-Dup. Between these two parts lies a part identical to both S1-Dup and LCR16a (in blue in panel C of Supplemental figure S8). We identified this overlapping region (207 bp) as the breakpoint of the duplication and confirmed an NAHR mechanism. This breakpoint region corresponds to an *AluJb*. Based on this, we defined the mechanism as *Alu*-mediated LCR16a-associated NAHR.

### 9. Tandem duplications (clusters 3-4) breakpoint analysis

A

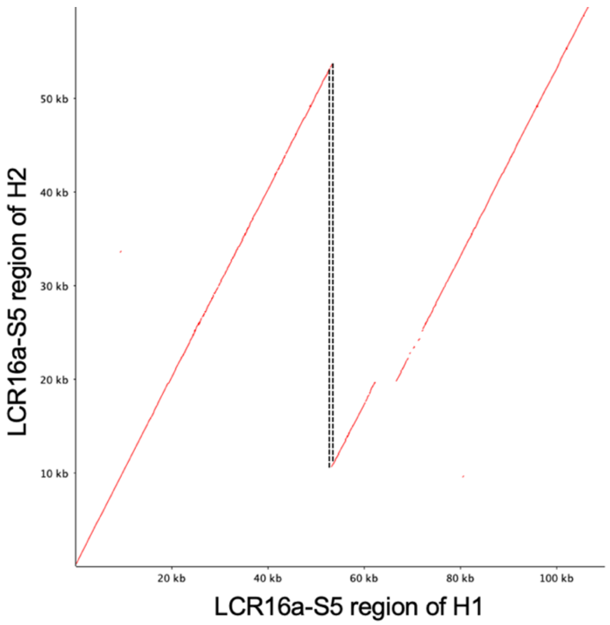


B

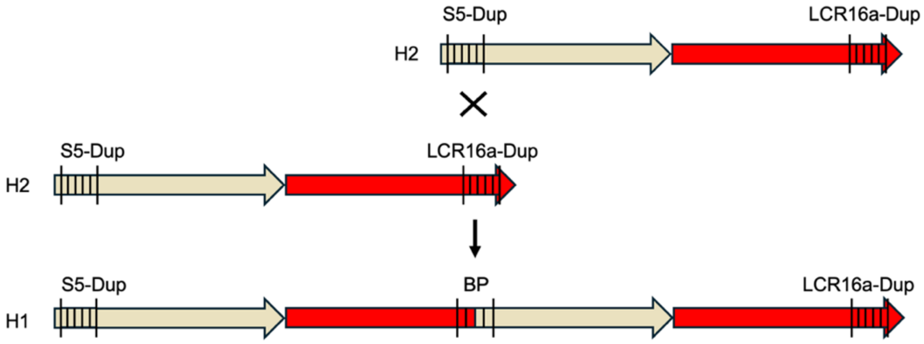


C

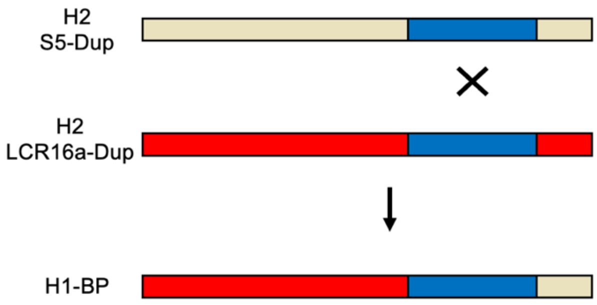


**Supplemental Figure S9.**

We identify the breakpoints of the tandem duplications in cluster 3 and 4 using a method similar to the one used to identify the breakpoints of the tandem duplications in cluster 1. Specifically, we used a dot-plot comparison between the S5-LCR16a segments of H2, which was considered as a reference, and the region containing all multiple copies of S5-LCR16a from the haplotypes where these duplications occur.

An example of a dot-plot comparing H2 S5-LCR16a and H1 (T2T) S5-LCR16a-S5-LCRC16 is shown in the figure. We observed that the start of S5 contains a sequence, named as S5-Dup, which is identical to a sequence found at the end of LCR16a, named as LCR16a-Dup (see figures A and B). In H1, this sequence was present in three copies: one at the start of S5, one at the end of LCR16a, and one in the region between the first LCR16a and the second S5, which was named BP. This pattern is consistent with duplications mediated by NAHR, where the mediating segmental duplication is typically present in three copies within the rearranged haplotype. The same pattern was observed across all haplotypes containing these tandem duplications.

To further investigate, we retrieved the sequences corresponding to S5-Dup, LCR16a-Dup, and BP from all the rearranged haplotypes, aligned them using MAFFT (Katoh et al. 2009), and manually inspected the alignments.

We found that the first part of H1-BP is more similar to H2 LCR16a-Dup, while the final part is more similar to H2 S1-Dup. Between these two parts, there is a segment identical to both S5-Dup and LCR16a (in blue in panel C). We identified this overlapping region (38 bp) as the breakpoint of the duplication and confirmed an NAHR mechanism. This region corresponds to an *AluSx*. Based on this, we defined the mechanism as *Alu*-mediated LCR16a-associated NAHR.
